# RNA G-quadruplex structures exist and function *in vivo*

**DOI:** 10.1101/839621

**Authors:** Xiaofei Yang, Jitender Cheema, Yueying Zhang, Hongjing Deng, Susan Duncan, Mubarak Ishaq Umar, Jieyu Zhao, Qi Liu, Xiaofeng Cao, Chun Kit Kwok, Yiliang Ding

## Abstract

Guanine-rich sequences are able to form complex RNA structures termed RNA G-quadruplexes *in vitro*. Because of their high stability, RNA G-quadruplexes are proposed to exist *in vivo* and are suggested to be associated with important biological relevance. However, there is a lack of direct evidence for RNA G-quadruplex formation in living cells. Therefore, it is unclear whether any purported functions are associated with the specific sequence content or the formation of an RNA G-quadruplex structure. Here, we profiled the landscape of those guanine-rich regions with the *in vitro* folding potential in the *Arabidopsis* transcriptome. We found a global enrichment of RNA G-quadruplexes with two G-quartets whereby the folding potential is strongly influenced by RNA secondary structures. Using *in vitro* and *in vivo* RNA chemical structure profiling, we determined that hundreds of RNA G-quadruplex structures are strongly folded in both *Arabidopsis* and rice, providing direct evidence of RNA G-quadruplex formation in living eukaryotic cells. Subsequent genetic and biochemical analysis showed that RNA G-quadruplex folding was sufficient to regulate translation and modulate plant growth. Our study reveals the existence of RNA G-quadruplex *in vivo*, and indicates that RNA G-quadruplex structures act as important regulators of plant development and growth.

## Introduction

The *in vivo* folding of RNA structure is tightly associated with its function and largely dependent on cellular context^1,2^. Because of the complexity within living cells, RNA folding *in vivo* could be very different from that *in vitro*^*3*,*4*^. For many complex RNA structures, although their folding *in vitro* has been well defined, the folding *in vivo* is poorly understood. One of such complex structures is RNA G-quadruplex (RG4), which is folded with guanine-rich (G-rich) sequences *in vitro* and consists of two or more layers of G-quartets involving both Hoogsteen and Watson-Crick base pairs^5,6^. RG4s can be very stable *in vitro* in the presence of ion cations such as potassium (K^+^), therefore they are hypothesized to exist *in vivo* and to be involved in novel functions^5^, such as post-transcriptional regulation of gene expression^7–9^. Nevertheless, the lack of direct evidences of RG4 folding *in vivo* raised the key question of whether all these suggested functions are due to RG4 structure or sequence motif. For example, a (GGC)_4_ motif was proposed to fold into an RG4 structure to repress translation of tumor-related genes^8^. However, without the evidence of *in vivo* folding, the translation inhibition may simply be due to the (GGC)_4_ sequence motif. Also, emerging evidences argues that this sequence motif is likely to form a stable hairpin RNA secondary structure rather than RG4^10^. Hence, it is crucial to determine whether RG4 truly exists *in vivo*, such that one can investigate and establish the relationship between RG4 and associated biological functions.

In recent decades, numerous efforts have been made to detect the folding of RG4s in fixed or living cells using G-quadruplex-specific antibodies^11^, ligands^12–14^ and fluorescent probes^15,16^. However, these methods suffer from three major shortcomings. Firstly, the antibodies / ligands / probes may induce the structure formation by perturbing the RNA G-quadruplex folding equilibrium in cells or binding to the G-rich sequence motifs^17,18^. Secondly, these methods cannot quantitively determine the folding state of individual G-rich regions of interest in cells. Thirdly, these methods cannot exclude the possibility of RG4 folding in fixing, permeabilizing or staining cells^17^. Because of these considerable shortcomings, these methods are considered inadequate for robustly determining the existence of RG4s and actual folding state of G-rich regions in living cells.

To address these shortcomings, two methods based on RNA chemical structure probing have been developed to determine folding state of G-rich regions in living cells. One direct method is based on the property of the chemical 2-methylnicotinic acid imidazolide (NAI) which preferentially modifies the last Gs of G-tracts when RG4 is folded. This special modification pattern is subsequently detectable by reverse transcription at both individual targeted RNAs and at a genome-wide scale^17,19^. The other method is more indirect, the modification is based on the specific methylation of the N7 position of guanine (N7G) by dimethyl sulfate (DMS)^20^. When a very high concentration of DMS (∼8%) is applied to the cells, all the N7 positions of G residues in the unfolded G-rich regions will be methylated *in vivo*^17^. These methylated G-rich regions are unable to refold into RG4 structure *in vitro* in the presence of K^+^. Since RG4 refolding *in vitro* are able to stall reverse transcriptase during reverse transcription (RT)^21^, *in vivo* unfolded G-rich regions are unable to lead to RT stalling^17^. On the opposite, if the G-rich regions are folded into RG4 structures *in vivo*, then N7G is protected from DMS modification. These unmodified G-rich regions are able to reform into RG4 structures later during RT, subsequently causing the RT stalling^17^. Both methods were applied in yeast and the DMS method was also applied in mouse embryotic stem cells (mESCs)^17^. These studies concluded that G-rich regions with the potential to form RG4s *in vitro* were globally unfolded *in vivo*. These results underpinned the fact that both yeast and mice avoid the formation of RG4s *in vivo*, and the presumption that RG4s do not exist in eukaryotic cells^17^.

Unlike yeast and animals, plants are sessile eukaryotic organisms and have evolved independently with unique regulatory strategies. For instance, cellular K^+^ is the most abundant ion in plants and plays key roles in plant development and stress response^22,23^. Given the importance of K^+^ in affecting RG4 folding, and the ability of plants in maintaining K^+^ balance within the cells, we hypothesize that plants are more likely to adopt RG4 structure *in vivo*. In addition, our previous study and others on individual G-rich sequences with folding potential *in vitro* have suggested that plants might favor RG4 structures^9,24^. However, the existence of RG4s *in vivo* in plants has not been determined and remains an open question.

Here, we investigated the *in vivo* folding state of G-rich regions in plants. Firstly, we profiled the landscape of the G-rich regions with the potential to fold into RG4s *in vitro* in the *Arabidopsis* transcriptome. We also revealed the unique RNA structural features of these regions. Using chemical structure probing, we found that those G-rich regions with *in vitro* folding potential are strongly folded in both *Arabidopsis* and rice. We further demonstrated that RG4 formation is sufficient to regulate gene expression, and subsequently plant growth. Taken together, these findings provided the first direct evidence of global RG4 formation in living eukaryotic cells, and revealed RG4s as important regulators for plant growth.

## Results

### Profiling of G-rich regions with potential to fold into RG4 structures in *Arabidopsis* transcriptome

To systematically search for G-rich regions with the potential to fold into RG4 structures in *Arabidopsis*, we used rG4-seq, an *in vitro* approach for detecting RG4 structures formation at a transcriptome-wide scale^25^. RG4 formation *in vitro* is stabilized by potassium ions (K^+^) (Fig. 1a), but not lithium ions (Li^+^), and is preferentially stabilized by G-quadruplex stabilizing ligands such as pyridostatin (PDS)^26^. G-rich regions which folded into RG4 structures *in vitro* can cause reverse transcriptional stalling (RTS)^21,25^. Therefore, RTS sites dependent on the presence of K^+^ or K^+^+PDS suggest the presence of G-rich regions with RG4 folding potential within sequences upstream of the RTS (Fig. 1b). We performed reverse transcription on *Arabidopsis* RNAs with Li^+^, K^+^ or K^+^+PDS respectively and generated corresponding libraries with high reproducibility (Supplementary information Fig. S1). To validate our rG4-seq, we mapped RT stops on the mRNA of *SUPPRESSOR OF MAX2 1-LIKE5* (*SMXL5)* which contains a G-rich region with RG4 folding potential identified recently^9^. We found a strong RT stalling at the 3’end of this G-rich region where the coverage dropped in the presence of both K^+^ and K^+^+PDS conditions, but not Li^+^ (Fig. 1c), agreeing with the previous gel-based assay^9^.

**Fig.1.**
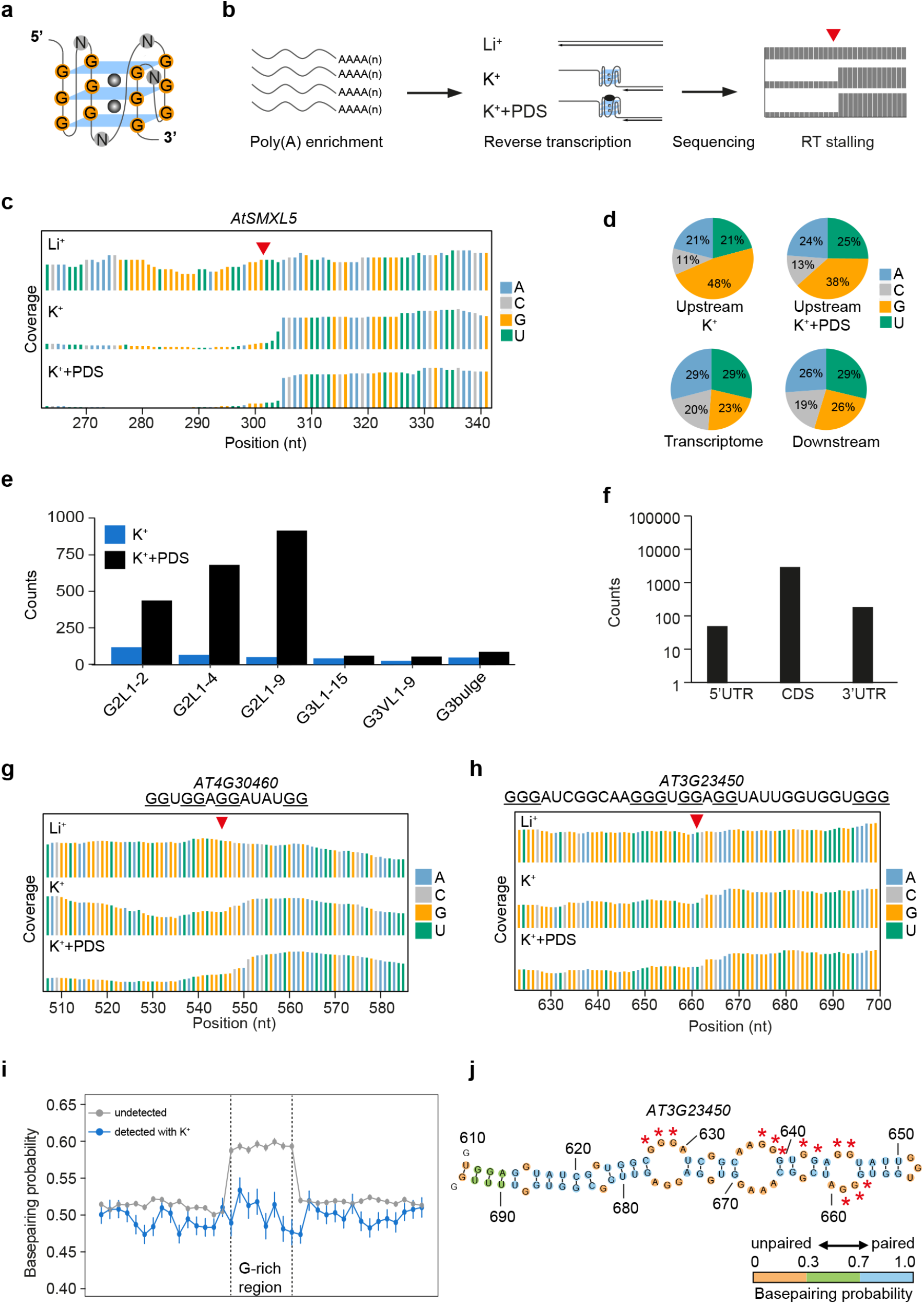
rG4-seq reveals the global landscape of G-rich regions with the potential to fold into RG4 structures in *Arabidopsis*. **a** Schematic of an RNA G-quadruplex (RG4). The schematic depicts an RG4 structure with three layers of G-quartets (G3 RG4, guanine (G) coloured in orange), with the loop length of any nucleotide (N, coloured in grey), potassium ions (K^+^, grey ball) coordinated within the G-quartets stabilize RG4s. **b** Workflow of rG4-seq. Poly(A) enriched RNAs were subjected to reverse transcription under the buffers with Li^+^ (non-stabilizing condition), K^+^ (stabilizing condition) or K^+^+PDS (stronger stabilizing condition), respectively. The G-rich region sites with folding potential were identified by comparing the coverage of reads between the rG4-seq libraries with different cations as described above. **c** rG4-seq profiles of *AtSMXL5* displayed the reads coverage of reverse transcription (RT) with Li^+^ (top), K^+^ (middle), and K^+^+PDS (bottom) respectively. The 3’end of the G-rich region is indicated by red triangle. A (blue), C (light grey), G(yellow), U(green). **d** Residue distribution around RTS sites. Guanine (G) was strongly enriched in the upstream sequences of RT stalling (RTS) identified under both K^+^ and K^+^+PDS conditions, but not in the transcriptome and the downstream sequences of RTS. A (blue), C (light grey), G(yellow), U(green). **e** Classification of G-rich regions with folding potential. G-rich regions with folding potential identified in K^+^ (dark blue) and K^+^+PDS conditions (black) were classified into six categories according to the number of G-quartets (G2 with two G-quartets or G3 with three G-quartets), loop length (L, 1-15 nt), and bulge size (non-canonical G3 RG4s with a guanine vacancy: G3VL1-9, or a bulge: G3bulge). **f** The prevalence of G-rich regions in different genic regions. **g** and **h** rG4-seq profiles of G2 G-rich region on *AT4G30460* (g) and G3 G-rich region on *AT3G23450* (h), otherwise in Fig. 1c. **i** Comparison of base-pairing probability (BPP) of alternative secondary structure in G-rich regions that are detected with K^+^(blue) and undetected (grey) using rG4-seq. The Gs in the G-rich region were classed to 8 bins, flanking sequences (100 nt on both sides) were classed to 20 bins, with 15 bins close to G-rich regions shown. Differences of BPPs between G-rich regions and flanking regions, detected by rG4 with K^+^, *P* = 0.444; undetected regions, *P* < 10^−16^; *P*-values, paired Student’s t-test. **j** Secondary structure of G-rich region detected by rG4-seq (in Fig. 1h) and flanking sequences on *AT3G23450*, predicted using *Vienna RNAfold*. The filling colours of orange, green and blue indicate the base-pairing probability of below 0.3, 0.3-0.7 and above 0.7 respectively. Red stars indicate the guanines comprising the G3 region. Numbers indicate nucleotide positions on the transcript.

We then searched for RTS sites dependent on the presence of K^+^ or K^+^+PDS in the *Arabidopsis* transcriptome. Our meta-analysis showed that guanine (G) was strongly enriched in sequences upstream but not downstream of RTS sites, for both K^+^ and K^+^+PDS conditions (Fig. 1d). A strong enrichment of guanine suggested the prevalence of G-rich regions with potential to form RG4 structures *in vitro*^17,25^. We searched for G-rich regions able to form RG4 structures on the basis of special sequence feature, GxLnGxLnGxLnGx (whereby G stands for Guanine; L stands for Loop, and x ≥2, n up to 15, see methods) in sequences upstream of RTS. In total, we found 381 and 2457 G-rich regions with strong RTS dependent on K^+^ and K^+^+PDS respectively (Supplementary information Table S1, Table S2). We then classified these G-rich regions according to G-quartet number, loop length and bulge size. In the presence of K^+^, we found 253 G2 (with two G-quartets) and 128 G3 (with three G-quartets) G-rich regions (Fig. 1e and Supplementary information, Table S1). In the presence of K^+^+PDS, we detected 2234 G2 and 223 G3 regions (Fig. 1e and Supplementary information, Table S2). As illustrated by rG4-seq profiles of individual regions on *AT4G30460* and *AT3G23450*, coverage dropped down in the presence of K^+^ and K^+^+PDS, but not Li^+^, at the 3’end of these G-rich regions, indicating folding of G2 and G3 RG4 structures (Fig. 1g, h). We also examined the location of these G-rich regions, and found that they were mostly localized in the coding regions (Fig. 1f, Supplementary information, Table S1 and Table S2). Taken together, we revealed the global *in vitro* landscape of G-rich regions with the potential to fold into RG4 structures in the *Arabidopsis* transcriptome.

### Features of *Arabidopsis* RG4 formable regions

Over 65,000 G-rich regions were predicted *in silico* to form RG4 structures in *Arabidopsis* based on the sequence feature of GxLnGxLnGxLnGx^6,27^. However, we detected less than 3000 regions by rG4-seq (Fig. 1e and Supplementary information, Table S1, Table S2), indicating most of predicted regions are unlikely to fold into RG4 structures *in vitro*. This prompted us to ask whether this low rate of *in vitro* folding is due to competition from alternative secondary structure formation, which might prevent RG4 formation across these predicted regions. To test this hypothesis, we used *Vienna RNAfold* software to predict the secondary structures of both *in silico* predicted but undetected G-rich regions and the detected RG4 regions by rG4-seq with K^+28^. We then calculated the base-pairing probability (BPP) of each nucleotide based on the predicted secondary structures and performed the meta-property analysis^28^. We found that, for the undetected G-rich regions, the BPPs of G-rich regions were significantly higher compared to flanking regions (Fig. 1i, P < 10^−16^, paired Student’s t-test), indicating these G-rich regions were folded into strong secondary structures. In contrast, for those detected G-rich regions in the presence of K^+^, no significant differences of BPPs were found between G-rich regions and flanking sequences (Fig. 1i, P = 0.444, paired Student’s t-test), and the BPPs of G-rich regions are strongly lower than that of the undetected regions (Fig. 1i, P < 10^−16^, Student’s t-test). Therefore, those regions detected *in vitro* by rG4-seq are likely to form weak secondary structure while the undetected regions are likely to fold into strong secondary structures (Fig. 1j and Supplementary information, Fig. S2). Taken together, our results indicate that alternative secondary structures in those G-rich regions strongly affect the potential of folding into RG4 structures *in vitro*.

### SHALiPE-Seq robustly determines the folding state of G-rich regions

Next, we asked if these G-rich regions with folding potential *in vitro* are able to fold into RG4 structures *in vivo*. Our previous studies have shown that both NAI and DMS are capable of penetrating plant cells^29,30^. We selected NAI rather than DMS for our *in vivo* determination of the folding state of the G-rich regions in plants, because the DMS method using high DMS concentration (8%), causes plant wilting as well as significant RNA decay^17,31^. To avoid unpredictable inaccuracies of structure determination implied by the high concentration of DMS method, we employed the NAI chemical probing method by coupling our previous method selective 2’-hydroxyl acylation with lithium ion-based primer extension (SHALiPE) with high throughput sequencing, which we termed SHALiPE-Seq^19,29^.

SHALiPE-Seq is based on the preferential modification of the last G in G tracts of folded RG4s by 2-methylnicotinic acid imidazolide (NAI) (Fig.2a)^17,19,32^. Strong modifications are able to cause reverse transcription stalling on these last guanines, which are detectable by deep sequencing. Significantly higher reads number can be found on these last guanines for folded G-rich regions, rather than unfolded ones (Fig. 2a). This method could discriminate the folded state from unfolded state of individual G-rich regions by showing reads number with or without the special pattern. Before applying this method to plants *in vivo*, we firstly established benchmark SHALiPE profiles for both folded and unfolded states of G-rich regions in plants *in vitro* in the presence of K^+^ (folded state) or Li^+^ (unfolded state), respectively (Fig. 2a, see methods). We assured our NAI modification efficiencies in the presence of K^+^ and Li^+^ were similar, as indicated by gel-based analysis on 18S rRNA (Supplementary information, Fig. S3a). We then generated the corresponding SHALiPE-Seq libraries with high reproducibility (Supplementary information, Fig. S3b, c).

**Fig.2.**
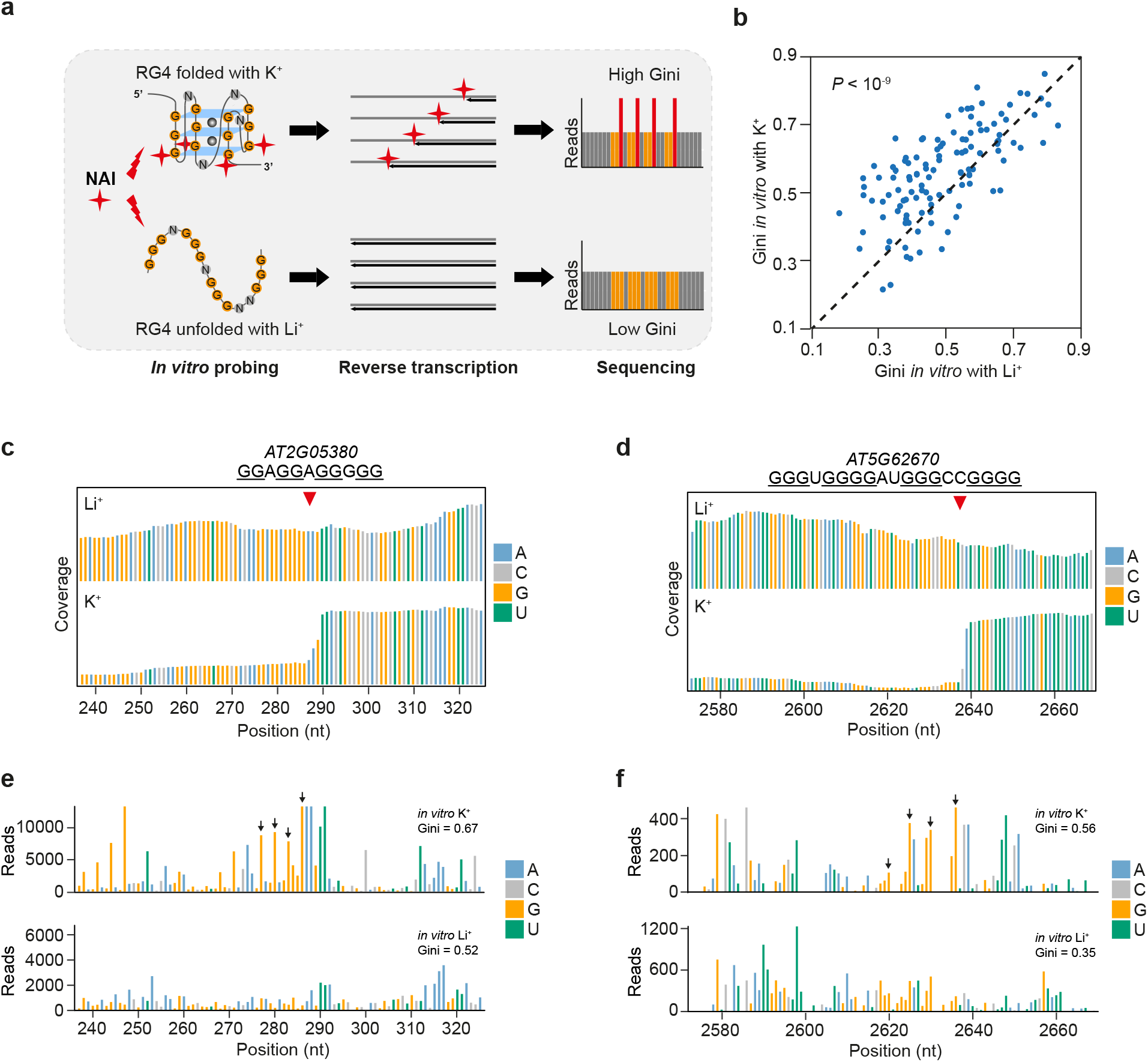
SHALiPE-Seq determines folding states of G-rich regions robustly. **a** Schematic of SHALiPE-Seq *in vitro* with K^+^ and Li^+^. *In vitro* probing in the presence of K^+^ and Li^+^ established the benchmarks of folded and unfolded states of G-rich regions respectively. NAI (indicated by red cross) preferentially modifies the last G in G tracts of folded RG4s, resulting in a high Gini index of read counts (reads on preferentially modified Gs were in red) in SHALiPE profiles. In contrast, the distribution of the SHALiPE profile for the unfolded state in the presence of Li^+^ is uniform, resulting in a low Gini. **b** For G-rich regions detected using rG4-seq in the presence of K^+^, Gini of SHALiPE profiles *in vitro* with K^+^ (folded state) was greater than that of *in vitro* with Li^+^ (unfolded state) by a factor of 1.20 (n = 117, *P*-value, paired Student’s t-test, average reads coverage on G ≥ 50 reads/nt). **c** and **d** rG4-seq profiles of G2 G-rich region on *AT5G05380* (c) and G3 G-rich region on *AT5G62670* (d). The 3’end of the G-rich region is indicated by a red triangle. A (blue), C (light grey), G(yellow), U(green). **e** and **f** SHALiPE profiles *in vitro* with K^+^ or Li^+^ of G2 G-rich region on *AT5G05380* (c) and G3 G-rich region on *AT5G62670* (d). High read counts of last guanines of G-tracts (indicated by black arrows) represent strong modifications of NAI on these guanines in the presence of K^+^, indicating RG4s are folded. In the presence of Li^+^, last Gs are not strongly modified, representing unfolded state of these G-rich regions. The Gini values with K^+^ are higher than those with Li^+^, as indicated. A (blue), C (light grey), G(yellow), U(green).

In these SHALiPE-Seq libraries, the distribution of read counts on guanines for the folded
state (with K^+^) is uneven, while the distribution for the unfolded state (with Li^+^) is uniform (Fig. 2a)^17^. These distributions were measured using the Gini index, where a high Gini indicates an uneven distribution (folded state) and a low Gini indicates a uniformed distribution (unfolded state) (Fig. 2a)^17^. As expected, for the G-rich regions detected by rG4-seq, the Gini of SHALiPE profiles *in vitro* with K^+^ was greater than that *in vitro* with Li^+^ by a factor of 1.20 (Fig. 2b, *P* < 10^−9^, paired Student’s t-test and Supplementary information, Table S3, reads ≥ 50 / nt). As illustrated by the individual G2 and G3 regions on *AT2G05380* and *AT5G62670* identified by rG4-seq (Fig. 2c, d), high reads counts were found on the last Gs (indicated with black arrows) in the SHALiPE profiles with K^+^, indicating the preferential NAI modification on these Gs when RG4s are formed (Fig. 2e, f, top channels). Conversely, reads counts in the SHALiPE profile with Li^+^ are more uniform, representing the unfolded state of these G-rich regions (Fig. 2e, f, bottom channels). As a result, the Ginis of the SHALiPE profiles in the presence of K^+^ (0.67 and 0.52) are higher compared to the corresponding 0.52 and 0.35 for Li^+^ (Fig. 2e, f, text annotation on upper right). These results indicate that SHALiPE-Seq and rG4-seq are in strong mutual agreement, and that SHALiPE-Seq profiling is able to determine the folding state of individual G-rich regions at the transcriptome-wide scale.

### Stable folding of RG4 structures in *Arabidopsis in vivo*

To assess the *in vivo* folding state of these G-rich regions, we firstly performed *in vivo* NAI chemical probing in *Arabidopsis*^29^, and generated *in vivo* SHALiPE-Seq libraries with high reproducibility (Supplementary information, Fig. S3d). We then compared SHALiPE profiles *in vivo* with our benchmark SHALiPE profiles *in vitro* for both folded and unfolded states (Fig. 3a)^17,19,29^. Next, we calculated the Gini *in vivo* of G-rich regions and found that Gini *in vivo* was greater than that *in vitro* with Li^+^ by a factor of 1.10 (Fig. 3b, *P* < 10^−16^, paired Student’s t-test and Supplementary information, Table S4, reads ≥ 50 / nt). This result suggested that these G-rich regions are folded into RG4 structures *in vivo*. To further quantify the folding state *in vivo*, we calculated the *in vivo* folding score: a comparison of Gini *in vivo* with Gini *in vitro* in the presence of Li^+^, scaled relative to Gini *in vitro* with K^+^ versus Gini *in vitro* with Li^+17^. *In vivo* folding scores of these G-rich regions centered near 1 with a median value of 0.755 (Fig. 3c and Supplementary information, Table S4), indicating that most G-rich regions identified by rG4-seq were strongly folded in *Arabidopsis*. The *in vivo* folding of RG4s in *Arabidopsis* differs from the unfolded states in yeast and mouse embryonic stem cells, where the folding scores centered near 0 (median values of −0.02 and 0.06 respectively)^17^. The *in vivo* SHALiPE profile resembled the *in vitro* SHALiPE profile in the presence of K^+^ (folded state), but not in the presence of Li^+^ (unfolded state) for individual G-rich regions, as exemplified by regions on *AT3G23450* and *AT4G30460* (Fig. 3d, e). We further assessed whether any specific type of G-rich regions may be preferentially folded *in vivo*, but found very similar folding scores among different types of RG4s indicating no specific preference (Fig. 3f). In addition, the folding scores for RG4s in both coding regions (CDS) and untranslated regions (UTRs) were quite similar (Fig. 3g), indicating both CDS and UTRs contain stable RG4 structures *in vivo*. Taken together, our results indicate that in *Arabidopsis*, hundreds of G-rich regions are strongly folded into RNA G-quadruplexes *in vivo*, unlike the previously reported *in vivo* observations in yeast and mice^17^.

**Fig.3.**
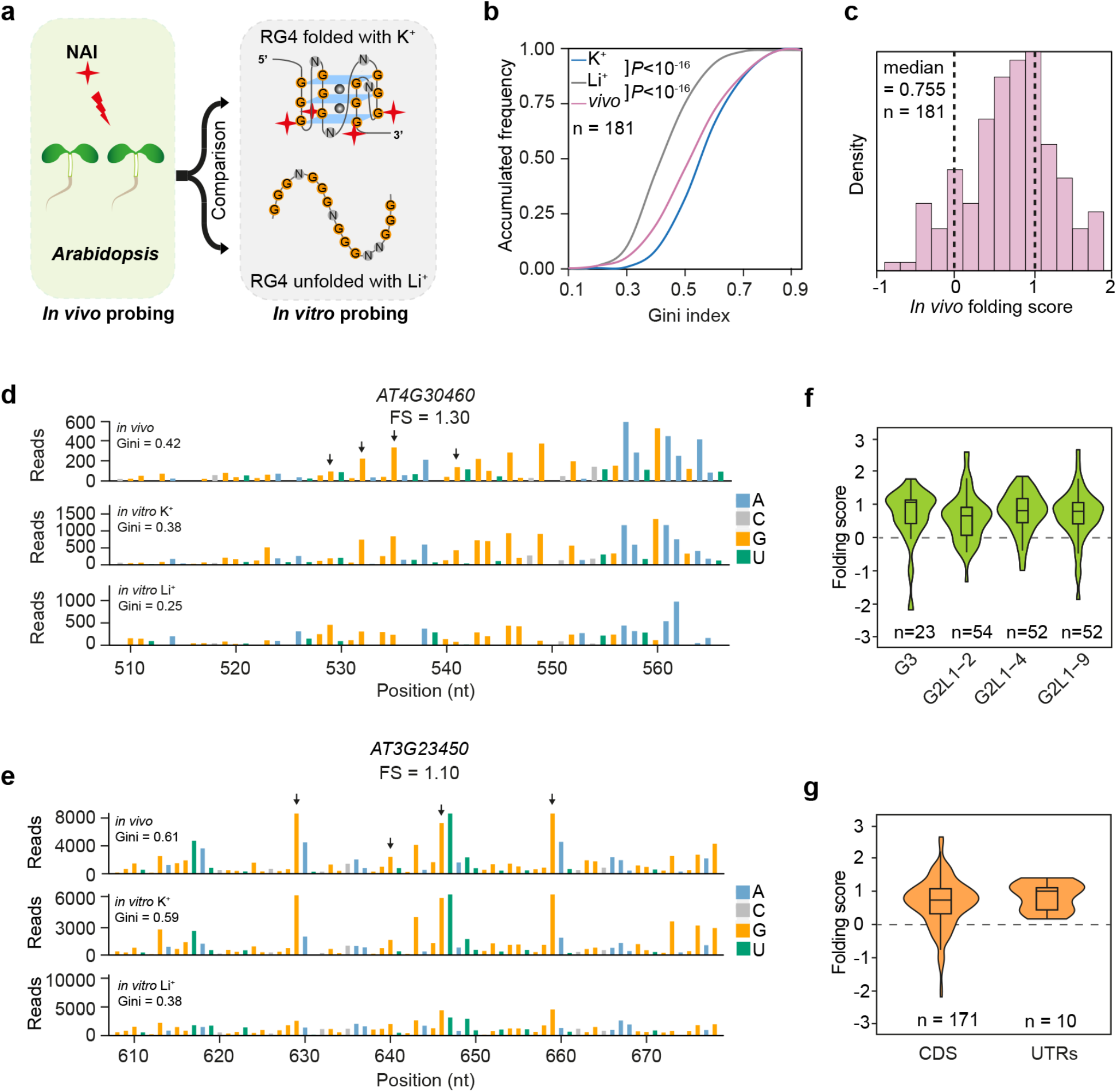
*In vivo* SHALiPE-Seq reveals hundreds of folded RG4s in *Arabidopsis.* **a** Schematic of *in vivo* SHALiPE-Seq in *Arabidopsis*. *In vivo* folding state of G-rich region is evaluated by comparing SHALiPE profiles *in vivo* with SHALiPE profiles *in vitro* in the presence of Li^+^ or K^+^ respectively. **b** Cumulative plot of Gini index of *Arabidopsis* G-rich regions *in vivo, in vitro* with Li^+^ and *in vitro* with K^+^. A significantly higher Gini *in vivo* than that *in vitro* with Li^+^ indicates the folding state of G-rich regions *in vivo*. *P*-value, paired Student’s t-test. **c** Histogram of *in vivo* folding score (FS) in *Arabidopsis.* The median value is 0.755. The FS of 0 represents the unfolded states (with Li^+^) and 1 represents the folded states of RG4s (with K^+^) *in vitro.* **d** and **e** SHALiPE profiles of the RG4 region on *AT4G30460* (d) and *AT3G2345*(e). The *in vivo* SHALiPE profile resembled the *in vitro* SHALiPE profile with K^+^ (the last Gs of G-tracts indicated by black arrows) but not the *in vitro* SHALiPE profile with Li^+^, indicating the folded state of this RG4 *in vivo*. **f** Violin plot of *in vivo* folding scores of G-rich regions of different types. Folding scores of RG4s of different catalogues are similar (*P* values > 0.05, one-way ANOVA/ Tukey HSD post hoc test). **g** Violin plot of *in vivo* folding scores of G-rich regions in CDS and UTRs. Folding scores of RG4s of different genic regions are similar (*P* > 0.05, one-way ANOVA/ Tukey HSD post hoc test).

### *In vivo* folding of RG4 structures in rice

To determine whether RG4s exist in diverse plant species, we investigated the folding state of G-rich regions in rice (*Oryza sativa* ssp. *japonica*), one of the most globally important crops^33^. We performed SHALiPE-Seq profiling in rice, and compared *in vivo* SHALiPE profiles with *in vitro* with K^+^ and Li^+^ respectively (Fig. 4a). Similar to observations in *Arabidopsis*, Gini *in vivo* for rice was greater than that *in vitro* with Li^+^ by a factor of 1.20 (Fig. 4b, *P* < 10^−16^, paired Student’s t-test and Supplementary information, Table S5). *In vivo* folding scores centered near 1 with a median value of 0.938, indicating the folding status of RG4s in rice (Fig. 4c and Supplementary information, Table S5). The *in vivo* SHALiPE profile resembled the *in vitro* SHALiPE profile in the presence of K^+^ (folded state) but not Li^+^ (unfolded state) for individual RG4s, as exemplified by regions on *LOC_Os02g15810* and *LOC_Os07g41694* (Fig. 4d, e). Moreover, no significant differences were found of the folding scores for RG4s of specific types nor different genic locations (Fig. 4f, g). Taken together, as represented by model dicot and monocot plant species, our results suggest the general existence of RG4s in the plant kingdom.

**Fig.4.**
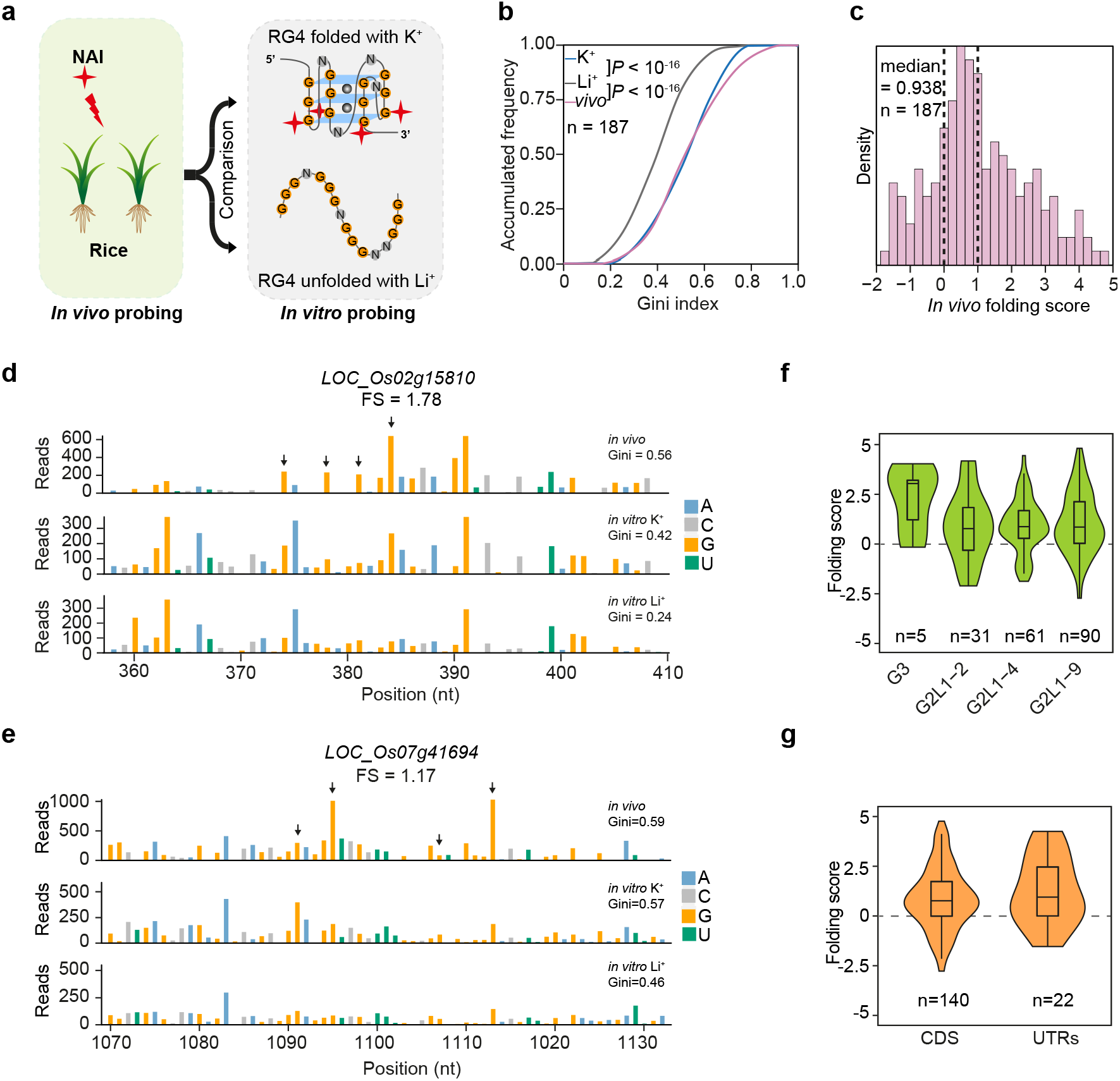
*In vivo* SHALiPE-Seq reveals hundreds of folded RG4s in rice. **a** Schematic of *in vivo* SHALiPE-Seq in rice. The *in vivo* folding state of G-rich region is evaluated by comparing SHALiPE profiles *in vivo* with SHALiPE profiles *in vitro* in the presence of Li^+^ or K^+^ respectively. **b** Cumulative plot of Gini index of rice G-rich regions *in vivo, in vitro* with Li^+^ and *in vitro* with K^+^. A significantly higher Gini *in vivo* than that *in vitro* with Li^+^ indicates the folding state of G-rich regions *in vivo*. *P*-value, paired Student’s t-test. **c** Histogram of *in vivo* folding score (FS) in rice. The median value is 0.938. The FS of 0 represents the unfolded states (with Li^+^) and 1 represents the folded states of RG4s (with K^+^) *in vitro.* **d** and **e** SHALiPE profiles of the RG4 region on *LOC_Os02g15810* (d) and *LOC_Os07g41694*(e). The *in vivo* SHALiPE profile resembled the *in vitro* SHALiPE profile with K^+^ (the last Gs of G-tracts indicated by black arrows) but not the *in vitro* SHALiPE profile with Li^+^, indicating the folded state of this RG4 *in vivo*. **f** Violin plot of *in vivo* folding scores of G-rich regions of different types. Folding scores of RG4s of different catalogues are similar (*P* values > 0.05, one-way ANOVA/ Tukey HSD post hoc test). **g** Violin plot of *in vivo* folding scores of G-rich regions in CDS and UTRs. Folding scores of RG4s of different genic regions are similar (*P* > 0.05, one-way ANOVA/ Tukey HSD post hoc test).

### RG4 structure regulates translation and plant development

Considering our demonstration of the prevalence of RG4s in plants, and that RG4s are hypothesized to be associated with post-transcriptional regulation of gene expression^5,6^. We wanted to test the hypothesis by examining whether RG4s function *in vivo*. We focused on those *in vivo* folded RG4s (folding score > 0.5) located in UTRs of genes in *Arabidopsis* (Supplementary information, Table S4). We screened T-DNA mutants of these genes and successfully identified homozygotes for gene *HIRD11*, which encodes a KS-type dehydrin and contains a G2 RG4 in its 3’UTR (Fig. 5a,b and Supplementary information, Fig. S4a). The mRNA abundance of *HIRD11* in the mutant (termed *hird11-1*) was strongly reduced compared to wildtype Col-0 (Fig. 5c). The growth of *hird11-1* was largely retarded relative to Col-0, represented by shorter primary roots compared to Col-0 (Fig. 5e, f). The phenotype of shorter primary roots in *hird11-1* was restored by complementing mutants with *HIRD11* containing wildtype RG4 sequence (wtRG4), indicating *HIRD11* promotes plant growth (Fig. 5d, e, f). Strikingly, when *hird11-1* mutant was complemented with *HIRD11* containing mutated RG4 sequence (mutRG4, G to A mutation to disrupt RG4 folding capacity, Fig. 5d), the primary root length of mutRG4 plants was distinctively longer than that of wtRG4 plants (Fig. 5e, f).

**Fig.5.**
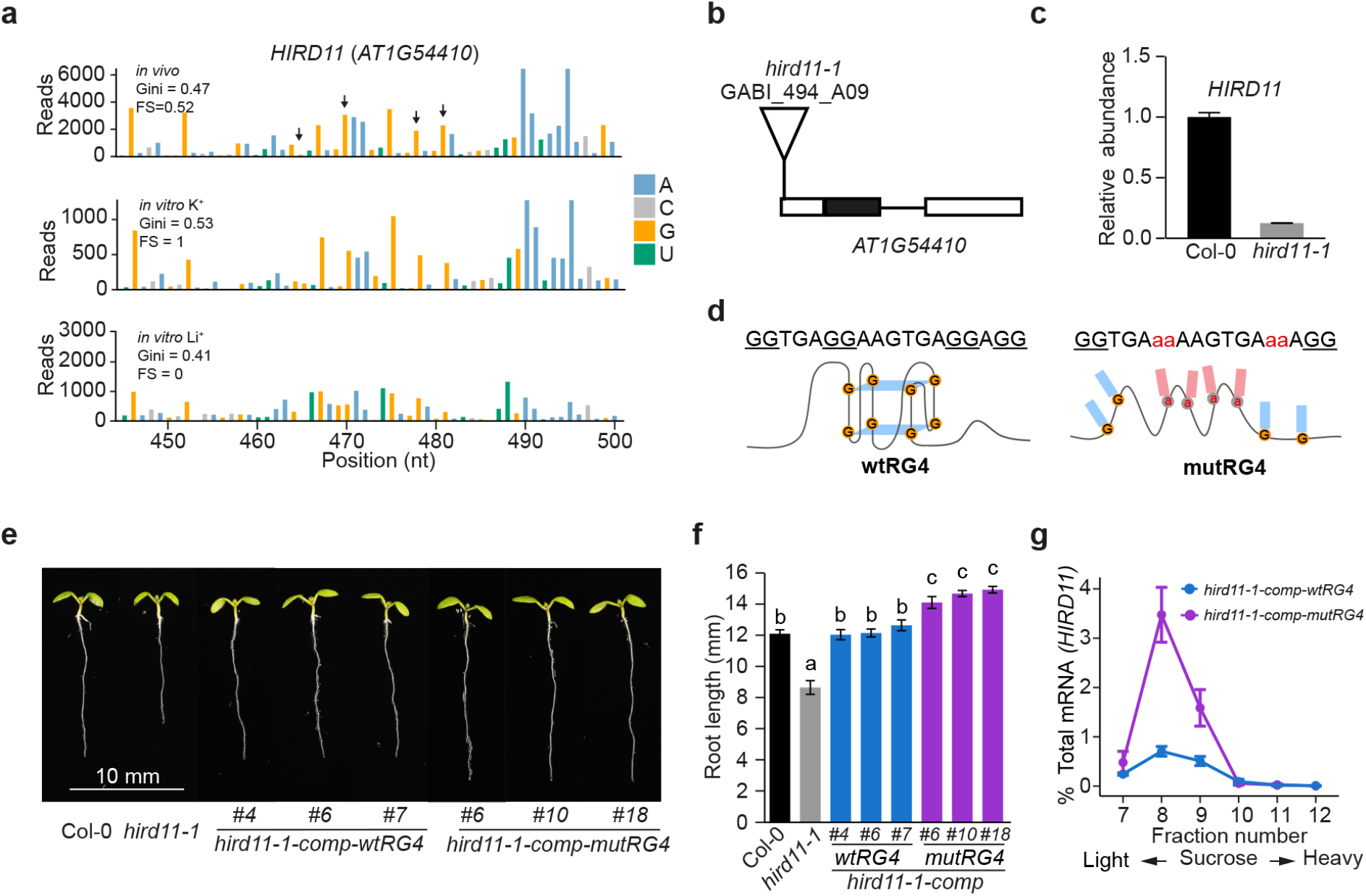
RG4 regulates plant growth and translation. **a** SHALiPE profiles of the RG4 region on 3’UTR of *HIRD11* (*AT1G54410*) for *in vivo*, *in vitro* with K^+^ and *in vitro* with Li^+^. The *in vivo* SHALiPE profile resembled the *in vitro* SHALiPE profile with K^+^ (the last G in G tracts of the RG4 region indicated by black arrows) but not the *in vitro* SHALiPE profile with Li^+^, indicating the folded state of this RG4 *in vivo*. **b** Schematic diagram of *HIRD11* showing the T-DNA insertion site of *hird11-1* (GABI_494_A09). **c** Relative mRNA abundance of *HIRD11* in Col-0 and *hird11-1* plants indicated the *hird11-1* is a knock-down mutant, error bars indicate SE. **d** Sequences of wildtype RG4 (wtRG4, left) and disrupted G-rich region with G to A mutation (mutRG4, right) on *HIRD11*. **e** and **f** RG4 modulates plant growth. Representative 6-day-old plants of Col-0, *hird11-1*, complemented *hird11-1* with wildtype RG4 (*hird11-1-comp-wtRG4*) and complemented *hird11-1* with mutated RG4 (*hird11-1-comp-mutRG4*) (e); and average primary root lengths (f) of more than 20 plants for each genotype. Significant differences were evaluated by one-way ANOVA/ Tukey HSD post hoc test (*P* < 0.05). Error bars indicate SE. **g** Analysis of polysome-associated *HIRD11* mRNA in the transgenic plants. RNA abundance of *HIRD11* in each polysome associated fraction was detected by qRT-PCR and quantified as a percentage relative to their total amount, error bars indicate SE.

We assessed whether RG4 folding affected the gene expression of *HIRD11*. Firstly, we compared the mRNA abundance of *HIRD11* in wtRG4 and mutRG4 plants, and found no significant difference of mRNA between these plants (Supplementary information, Fig. S4b). We then determined whether RG4 folding may affect *HIRD11* translation. Since no commercial antibody against HIRD11 is available, we used polysome analysis by combining sucrose gradient fragmentation with qRT-PCR to detect the translational level of *HIRD11* in these plants^34,35^. We found that polysome associated *HIRD11* mRNA in mutRG4 plants was much higher than that in wtRG4 plants (Fig. 5g), indicating a higher translational level of HIRD11 in mutRG4 plants. Therefore, our results indicate the RG4 on *HIRD11* suppresses its translation to modulate plant growth and development.

## Discussion

‘If G-quadruplexes form so readily *in vitro*, nature will have found a way of using them *in vivo*’ (Statement by Sir Aaron Klug over 30 years ago)^36^. Following decades of research by the community exploring RG4 structure across living eukaryotic cells, here for the first time, we determined hundreds of RG4s folded in model dicot and monocot plant species, providing direct evidence of RG4 existence in eukaryotes (Figs. 3 and 4). By both genetic and biochemical validation, we also demonstrated the important roles of the RG4 structure in modulating plant growth and development (Fig. 5).

RG4 structures contains unique sequence features of GxLnGxLnGxLnGx (whereby G stands for Guanine; L stands for Loop, and × ≥2, n up to 15). For decades, *in silico* prediction based on sequence features was used to search for putative RG4s at the genome-wide scale in different organisms^6,27,37,38^. With the rise of deep sequencing methods, high throughput methods such as rG4-seq were developed to map G-rich regions with *in vitro* RG4 folding potential throughout the transcriptome^25^. Here, we performed rG4-seq to map these G-rich regions in the *Arabidopsis* transcriptome. We identified less than 3000 G-rich regions with RG4 folding potential *in vitro* (Supplementary information Table S1, Table S2). In contrast, the *in silico* sequence-based method predicted over 65,000 G-rich regions^27^. Following our assessment of RNA secondary structures, we found that alternative secondary structures within those G-rich regions folded into might compete with RG4 structure formation (Fig. 1i and j). If the G-rich region is able to form strong secondary structure, it is unlikely to fold into RG4 structure. This might be due to the slower folding kinetics for RG4 structure formation compared to the formation of alternative secondary structures^39^. Thus, from a transcriptome-wide perspective, RG4 folding capability is generally influenced by alternative secondary structure formation, which may be an important way of regulating RG4 folding potential.

Following our rG4-seq *in vitro* profiling, we also revealed unique features for G-rich regions with RG4 folding potential in plants. We found that G-rich regions with potential to form G2 RG4s rather than G3 RG4s are predominate (>90%) in plants (Fig. 1e, Supplementary information Table S1, Table S2), whereas in human cells, G-rich regions with potential to fold G3 RG4s are dominant^25^. Rather than the very stable structures that G3 RG4s fold (Tm >55 °C), G2 RG4s are less stable (Tm ∼ 14-30 °C)^40,41^, thereby harbouring higher flexibility to switch between folded and unfolded states within the temperature range that most plants favour. This flexibility may facilitate a regulatory role in plant adaption to the immediate local environment. Rather than the depletion of G-regions for folding into RG4 structure in bacteria or avoiding formation of stable G3 RG4s in animals^17^, plants seem to have evolved a preference for G2 RG4 formation, which might be one of the reasons why plants have adopted RG4 structure as important regulators.

The large number of RG4s present in plants (Figs. 3 and 4) is probably explained by the physiological conditions in plants being favourable towards RG4 structure formation. For example, potassium (K^+^) is the predominant cytoplasmic inorganic cation in plants with a concentration around 100 mM^42,43^, which is preferable for RG4 folding. Notably, unlike animals, plants lack a potassium/sodium exchanger and thus use a unique accumulation and release system for potassium^43^. Since RG4 folding is highly sensitive to potassium levels, RG4 structures have great potential to be adopted as a regulator in response to dynamic potassium level changes. Our gene ontology analysis (Supplementary information, Fig. S5) confirmed that genes involved in metal ion binding are significantly enriched (Fig. S5). As such, RG4 structures might contribute to maintaining an optimum potassium level in plants. Apart from potassium levels, temperature is another key factor that was suggested to affect the folding of RG4s *in vitro*^40,41^. The optimal environmental temperature for plants (21-22 °C for *Arabidopsis* and 26-28 °C for rice) is much lower than the body temperature of animals (37 °C for human and 36.6 °C for mice). Thus, this relatively low temperature may allow the stable formation of RG4 structures in plant cells. In addition, a recent study found that the *Arabidopsis* RNA binding protein, zinc-finger protein JULG1 preferentially binds to a specific G-rich sequence with *in vitro* folding potential, stabilizing the RG4 structure *in vitro* even in the absence of potassium^9^. This result indicated that co-factors such as RNA binding proteins might be important regulators for the formation of RG4 structures in plants. Notably, *in vivo* folding scores of many individual regions (60 out of 181 in *Arabidopsis*, ∼1/3) are over 1 (Fig. 3c and Supplementary information, Table S4), indicating the *in vivo* folding state could be stronger than the *in vitro* folding state in the presence of K^+^. This result suggested that other factors might also promote RG4 formation *in vivo*. Hence, the ideal physiological conditions in plants together with those co-factors may confer plants with the ability to adopt RG4 structures, as regulators for gene expression.

Another interesting observation from our study is the variable folding states between *Arabidopsis* and rice (Fig. 3c and Fig. 4c). The distribution of *in vivo* RG4 folding scores in rice was shifted more to 1 compared to *Arabidopsis* (Fig. 3c and Fig. 4c). More RG4s in rice showed folding scores over 1 compared to the RG4s in *Arabidopsis* (Table S4 and S5). This result suggests that the folding landscape of RG4s is likely to be unique in different organisms under different growth conditions. Although previous chemical profiling in animal cell lines showed that RG4 structures are globally unfolded^17^, this might be due to the certain cell types having conditions not favourable for RG4 formation. Given the suggested functions of RG4s in special biological relevance, such as cancer cell growth and neurodegenerative diseases^7,8,18,44,45^, RG4 structures might stably form in specific cell types and growth conditions.

Previously, without direct evidence for RG4 formation in eukaryotes, it was not possible to infer whether suggested functions such as translation inhibition and splicing regulation are due to RG4 formation or specific sequence content^8,9,46,47^. In our study, we selected an *in vivo* folded RG4 located on *HIRD11* to assess the functional impact of this RG4 structure (Fig. 5). The compelling phenotypic difference in root length between plants with and without RG4 structure indicated the significant impact of RG4 structure on plant development (Fig. 5). The direct evidence of *in vivo* RG4 formation substantiated by both genetic and biochemical validations provides the first demonstration of RG4 structure functionality *in vivo*. Given the large number of RG4s present in plants (Figs. 3 and 4), further research is warranted to demonstrate that RG4s could significantly influence many aspects of plant growth and development. Apart from the translational regulation we revealed here (Fig.5), other biological functions such as splicing regulation might be also associated with RG4 structures^47^. Further exploration of the diverse functional roles of RG4 structures will expand our knowledge of RG4-dependent regulation of gene expression.

## Acknowledgments

We thank Prof. Giles Oldroyd (SLCU, Cambridge) and Dame Prof. Caroline Dean (John Innes Centre), Sir Prof. Shankar Balasubramanian (U. Cambridge), Prof. Alison Smith (John Innes Centre) and Dr. Desmond Bradley (John Innes Centre) for discussions with this work. This work was supported by the Biotechnology and Biological Sciences Research Council (BBSRC: BB/L025000/1 and BB/N022572/1) and the European Research Council (ERC: 680324) to Y.D.. M.I.U., J.Z., and C.K.K. were supported by Research Grants Council of the Hong Kong SAR (CityU11100218, N_CityU110/17, CityU 21302317), Croucher Foundation (9500030, 9500039), City University of Hong Kong (9680261), and the Petroleum Technology Development Fund (Nigerian government). X.C. was supported by the National Natural Science Foundation of China (31788103/91540203) and the Chinese Academy of Sciences (QYZDY-SSW-SMC022).

## Author contributions

X.Y. and Y.D. conceived the study; X.Y., C.K.K. and Y.D. designed the study; X.Y., Y.Z., H.D., S.D., M.I.U., J.Z., Q.L., C.K.K. and Y.D. performed the experiments; X.Y. and J.C. did the analyses; X.C., C.K.K., and Y.D. supervised the analyses; X.Y., and Y.D. wrote the paper with input from all authors.

